# NF-κB/p52 augments ETS1 binding genome-wide to promote glioma progression

**DOI:** 10.1101/2023.01.11.523507

**Authors:** Nicholas Sim, Li Yinghui

## Abstract

Gliomas are among the most invasive and chemo-resistant cancers, making them challenging to treat. Chronic inflammation is one of the key drivers of glioma progression as it promotes the aberrant activation of inflammatory pathways such as NF-κB signalling which drives cancer cell invasion, angiogenesis and tissue remodelling. NF-κB factors typically dimerize with its own family members, but emerging evidence of their promiscuous interactions with other oncogenic factors have been reported to activate the transcription of new target genes and function. Here, we show that non-canonical NF-κB activation directly regulates p52 at the *ETS1* promoter to activate its expression. This in turn impacts the genomic and transcriptional landscape of ETS1 in a glioma-specific manner. We further show that enhanced non-canonical NF-κB signalling promotes the co-localization of p52 and ETS1, resulting in the transcriptional activation of non-κB and/or non-ETS glioma-promoting genes. We conclude that p52-induced ETS1 overexpression in glioma cells remodels the genome-wide regulatory network of p52 and ETS1 to transcriptionally drive cancer progression.

## Introduction

Gliomas are one of the deadliest cancers and account for roughly 80% of all central nervous system (CNS) malignancies (1). A well-known driver and hallmark of glioma progression is chronic inflammation, which manifests through the infiltration of tumor-associated macrophages, upregulation of inflammatory cytokines, angiogenesis and tissue remodelling (2, 3). As levels of inflammatory mediators such as TNFα and IL-1β are elevated, it causes the aberrant activation of the nuclear factor kappa-light-chain-enhancer of activated B cells (NF-κB), mitogen activated protein kinase (MAPK)/extracellular-signal-regulated kinase (ERK) and signal transducer and activator of transcription 3 (STAT3) pathways which promote glioma progression (3, 4). The NF-κB signalling pathway, consists of the canonical, non-canonical, atypical and the DNA damage pathway, and is essential for regulation of multiple immune, inflammatory and developmental responses in normal tissues (5). When hyperactivated, it can promote the development and progression of various cancers including gliomas (6), breast cancers (7) and lymphomas (8). Among these pathways, the canonical and non-canonical signalling arms are the most well-studied. Canonical NF-κB signalling can be activated through various cytokines, growth factors or pattern-recognition receptors (9). Upon activation, the IκB kinase (IKK) complex is phosphorylated which in turn phosphorylates IκBα, signalling it for ubiquitin-dependent proteasomal degradation. This results in the nuclear translocation of p50/p65 dimers which can either activate or suppress gene expression (10). Conversely, non-canonical NF-κB signalling is triggered by a different subset of ligands such as B-cell activating factor (BAFF), receptor activator of nuclear factor kappa-? ligand (RANKL) and tumour necrosis factor-like weak inducer of apoptosis (TWEAK) (5, 6). This promotes the stabilisation of NF-κB-inducing kinase (NIK), which activates IKKα and thereby leads to the proteolytical processing of p100 into p52 (5). The resultant mature p52 then translocates into the nucleus as p52/RelB dimers to exert transcriptional regulation.

Another key mediator of inflammation is the E26 (ETS) transcription factor (TF) family which are transcriptionally regulated downstream of MAPK signalling (11). ETS TFs are involved in a wide range of biological processes which include cell cycle control, cell proliferation, angiogenesis, tissue remodelling and differentiation (12, 13). ETS factors such as ETV2 and PU.1 have been reported to be overexpressed in gliomas and their dysregulation promotes tumorigenesis and metastasis (14).

Traditionally, NF-κB subunits form specific homo- or hetero-dimers among themselves to exert diverse transcriptional functions. However, various TFs such as STAT3, p53 and ERG have also been demonstrated to associate with NF-κB to repress or enhance the transcriptional activation of κB-dependent genes in human cancers (15–17). Notably, NF-κB/p52 has been shown to physically interact and cooperate with ETS1 to regulate telomerase reactivation via the mutant telomerase reverse transcriptase (*TERT*) promoter, which is a cancer driver gene in almost 90% of human cancers (6, 18). However, the genome-wide regulatory interplay between p52 and ETS1 has yet to be further explored. Here, we show that p52 activation directly regulates *ETS1* expression through p52 binding at its promoter. The p52-driven overexpression of ETS1 resulted in the alteration of its genome-wide binding dynamics, which induced a change in the enrichment of its associated motifs and target genes. We further demonstrate that ETS1 functionally cooperates with p52 to promote glioma invasion and cell proliferation. Altogether, our data indicate the crucial role of NF-κB/p52 activation in enhancing the genomic binding landscape of ETS1 to drive glioma progression.

## Results

### NF-κB/p52 activation alters the genomic binding landscape of ETS1

The non-canonical NF-κB pathway can be activated in response to a variety of ligands including TWEAK (19). To investigate the role of NF-κB/p52 in regulating the genomic occupancy of ETS1 in gliomas, we performed chromatin immunoprecipitation (ChIP) sequencing (ChIP-seq) using p52 and ETS1 antibodies in the glioma cell line, U-87 MG, with and without TWEAK treatment. There were 270 and 3697 unique p52 and ETS1 binding sites obtained respectively in untreated cells. Following TWEAK treatment, 11503 p52 and 3852 ETS1 binding sites were identified. (Supplementary Fig. 1). Interestingly, we detected the enrichment of p52 at the *ETS1* promoter upon TWEAK stimulation (Fig. 1a). Consequently, TWEAK-induced p52 activation resulted in the upregulation of ETS1 expression (Fig. 1b). We further confirmed this effect through CRISPR-mediated knockdown (KD) of NFKB2 which inhibited TWEAK-induced upregulated of ETS1 (Supplementary Fig. 2).

**Figure 1.**
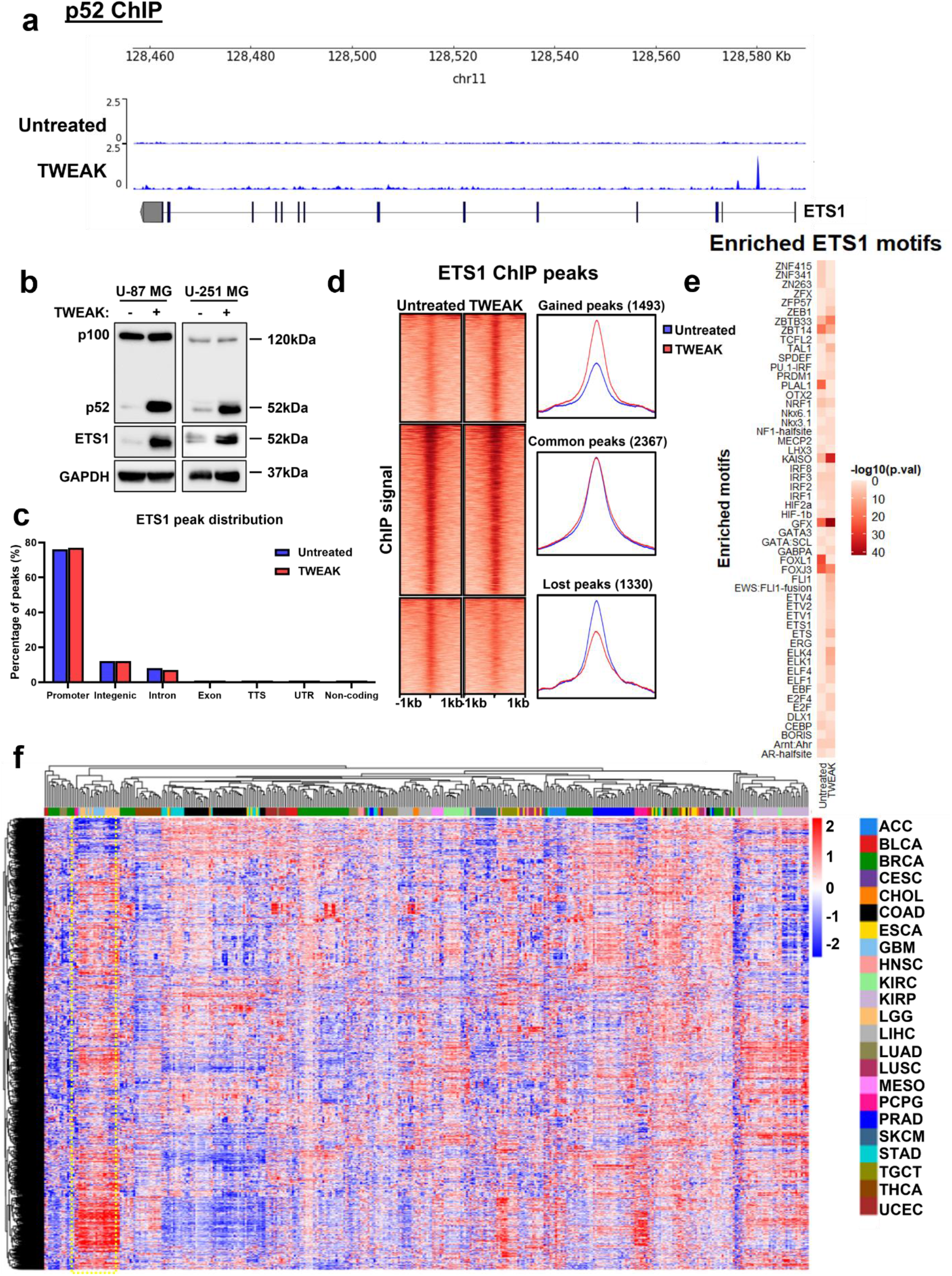
p52 activation remodels ETS1 DNA binding dynamics in glioma cells. (a) Genome browser view of *ETS1* gene locus and p52 ChIP-seq tracks in untreated and TWEAK-treated U-87 MG cells. Following TWEAK activation, p52 binds to the *ETS1* promoter. (b) ETS1 protein expression analysed by western blotting when NF-κB/p52 is activated in glioma cell lines. (c) Genomic distribution of ETS1 DNA binding peaks in untreated and TWEAK-treated U-87 MG cells. (d) DNA binding dynamics of ETS1 in untreated and TWEAK-treated U-87 MG cells. Signal intensity across each region (± 1 kb) is represented as a heatmap and the average log2 fold change in signal of gained, lost or common ETS1 peaks following TWEAK treatment is displayed at the right. (e) Heatmap representation of the enriched ETS1 motifs in U-87 MG cells with and without TWEAK-induced NF-κB/p52 activation. (f) Heatmap depicting the chromatin accessibility signal at ETS1 regions following TWEAK activation in TCGA Pan-cancer ATAC-seq dataset. ATAC-seq signals of glioma patient samples (LGG – Brain Lower Grade Glioma, GBM – Glioblastoma multiforme) are marked in yellow dotted box. TCGA study abbreviations can be found in https://gdc.cancer.gov/resources-tcga-users/tcga-code-tables/tcga-study-abbreviations.

Although p52 activation elevated ETS1 levels, it had little effect in both the number and distribution of ETS1 ChIP-seq peaks (Supplementary Fig. 1, Fig. 1c). However, we observed a distinct shift in the genomic binding landscape of ETS1 following TWEAK treatment (Fig. 1d). Further analysis of these differential sites revealed the increased enrichment of motifs for other ETS TFs such as FLI1, ELK4 and ELK1 (Fig. 1e), which have been implicated in cancer progression, following TWEAK-induced p52 activation (20–22). These TFs were also among the most highly expressed ETS factors in glioma patients (Supplementary Fig. 3). We also observed the enrichment of motifs for non-ETS factors, including KAISO and ZEB1, that have been documented to mediate the tumorigenic progression of gliomas (23, 24). This suggested that the p52-driven augmentation of ETS1 genomic binding may increase the co-association and cooperativity between ETS1 and different transcriptional co-activators that are overexpressed in gliomas. These changes in the ETS1 DNA binding dynamics may potentially enhance the transcription of various new target genes which can promote cancer progression.

To gain molecular insights into the relevance of the TWEAK-induced ETS1 genomic profiles in gene regulation, we interrogated the TCGA pan-cancer ATAC-seq dataset (25). Chromatin accessibility promotes TF occupancy and the activity of DNA regulatory elements which is crucial for gene transcription. When dysregulated in cancers, it can result in the overexpression of oncogenes through remodelling of enhancer signatures (26), pioneer factor-driven *de novo* enhancer formation (27) and/or chromosomal topology disruption (28). Among the 23 cancers within the TCGA ATAC-seq dataset, we unexpectedly found that the regions of accessible DNA which overlapped p52-activated ETS1 binding sites were highly enriched and specific to glioma patients (Fig. 1f). Collectively, these findings demonstrate that p52 activation drives a glioma-specific alteration in the genomic landscape of ETS1 that is enriched in accessible chromatin. Such DNA binding dynamics can promote the co-association of ETS1 with other oncogenic TFs to drive cancer-specific gene transcription.

### p52-activated ETS1 alters the transcriptomic landscape in glioma

To investigate the transcriptional regulatory role of TWEAK-induced p52 activation in glioma, we performed ChIP-seq using RNA polymerase II (RNA pol2) antibody in U87-MG treated with and without TWEAK. Differential analysis revealed that p52 activation alters genome-wide RNA pol2 recruitment where 412 sites are significantly upregulated, and 456 sites are significantly downregulated. Interestingly, we find that ETS1 is involved in roughly 20% of all upregulated RNA pol2 sites (Fig. 2a). We further verified the requirement of ETS1 for the maintenance of these upregulated RNA pol2 regions through *NFKB2* and *ETS1* KD which abolished TWEAK-driven RNA pol2 recruitment at these sites (Fig. 2b). These results show that p52-induced ETS1 is involved in the recruitment of transcriptional machinery.

**Figure 2.**
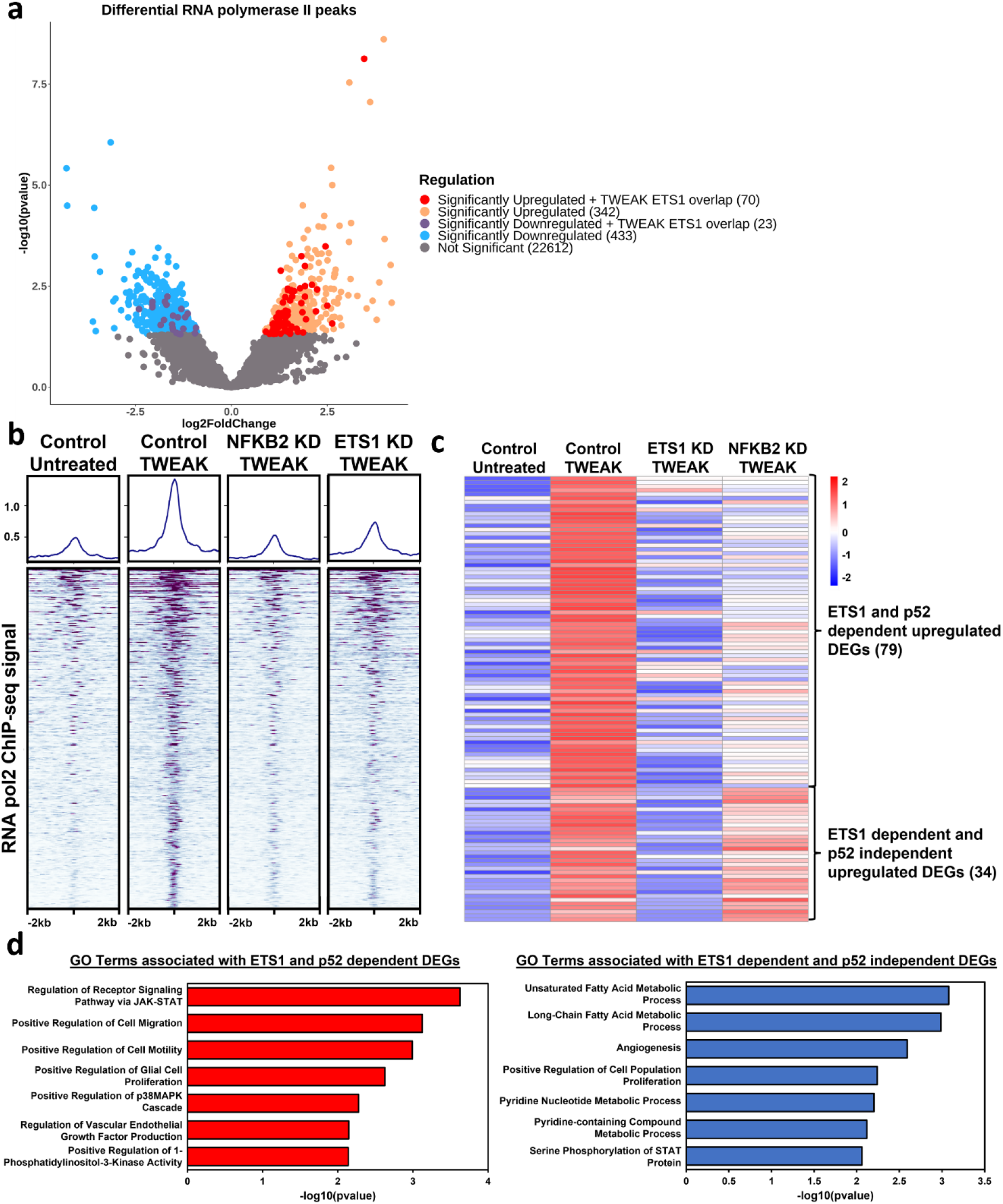
p52-driven ETS1 alters glioma RNA polymerase II recruitment and transcriptomic landscape. (a) Volcano plot depicting differential RNA polymerase II binding following NFκB/p52 activation and those bound by ETS1. (b) DNA binding heatmap of the significantly upregulated RNA polymerase II sites following NFκB/p52 activation and the effect of NFKB2 and ETS1 KD. (c) Heatmap depicting ETS1-dependent significantly upregulated differentially regulated genes following NFκB/p52 activation. (d) Bar chart depicting enriched GO terms from significantly upregulated ETS1 and p52 dependent DEGs as well as ETS1 dependent and p52 independent DEGs.

Following this, we performed RNA-seq to further to investigate the functional roles of p52-activated ETS1. This revealed that TWEAK treatment led to upregulation of 113 ETS1-dependent differentially expressed genes (DEGs). Among which, 79 DEGs were also dependent on p52, indicating that these genes are driven by p52-induced ETS1, while 34 DEGs were upregulated independent of p52 (Fig. 2c). Next, we performed gene ontological analysis on these two groups of upregulated DEGs. This analysis showed that both groups of DEGs shared enrichment of similar biological processes relating to proliferation and STAT activation. However, we were intrigued that the ETS1 and p52 dependent DEGs contained specific enrichment for terms associated with cell migration, VEGF production and MAPK activation (Fig. 2d). Taken together, these findings demonstrate the role of p52-induced ETS1 genomic binding in the transcriptional activation of distinct gene signatures that are associated with cancer-promoting processes.

### p52-induced ETS1 expression drives glioma invasion and cell proliferation

To functionally validate the role of p52-activated ETS1 in glioma progression, we targeted *NFKB2* and *ETS1* expression in U-87 MG and U-251 MG cells using CRISPR-Cas9 gene editing tools and verified their downregulation through western blotting (Supplementary Fig. 2,4). Here, we found that TWEAK-induced non-canonical NF-κB activation resulted in the increased proliferation and invasion of both glioma cell lines (Fig. 3a,b). However, these phenotypes were abolished following the loss of NFKB2 or ETS1 expression (Fig. 3a,b). These observations demonstrate that TWEAK-induced p52 activation promotes glioma progression through ETS1 and suggest that the gene regulatory roles of the p52-ETS1 interplay may be glioma specific.

**Figure 3.**
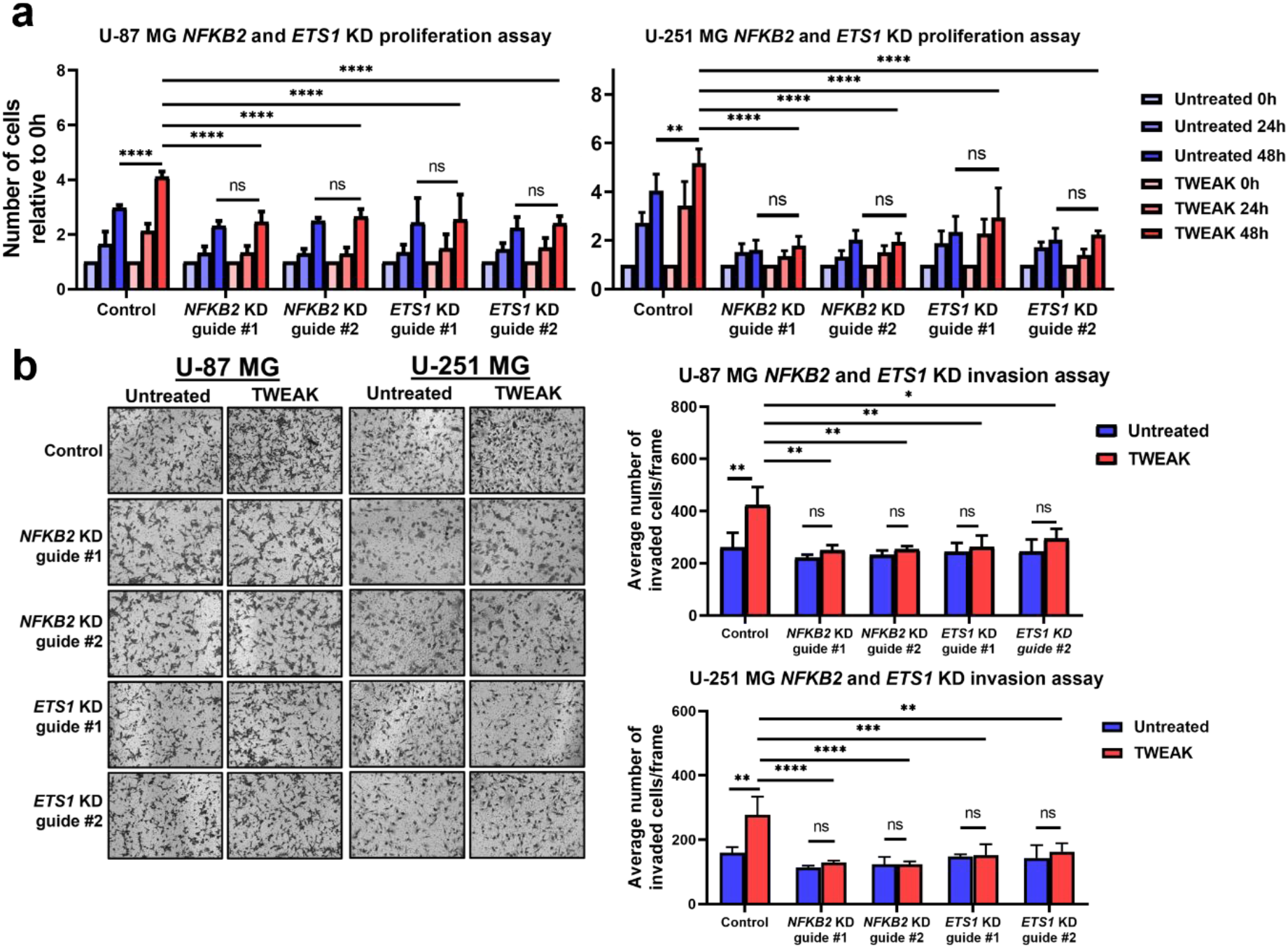
p52 drives the invasion and cell proliferation of glioma cells through ETS1. (a) Proliferation assay plot depicts the average number of cells counted relative to day 1 (mean ± s.d) in U-87 MG and U-251 MG *ETS1* KD cells with and without TWEAK-induced NF-κB/p52 activation. Data shown represent *n*=3 biological replicates. (b) Transwell invasion assay was performed in U-87 MG and U-251 MG *ETS1* KD cells with and without TWEAK treatment. Representative images from *n*=3 biological replicates are shown. Representative plots depict the average number of invaded cells (mean ± s.d) across five fields. Two-way ANOVA was used for all statistical analysis where \**P* < 0.05; \*\**P* < 0.01; \*\*\**P* < 0.001.

### ETS1 is crucial for the maintenance of p52 DNA binding

ETS1 can physically interact with various TFs such as Runx1, Pit-1 and HIF-2α where the DNA binding affinity of ETS1 or its binding partner are modulated (29–31). As ETS1 also physically co-associates and cooperates with p52 in gliomas (6, 18), we sought to examine the role of p52-activated ETS1 expression in the genomic binding of p52. To investigate this, p52 ChIP-seq was performed on CRISPR control and *ETS1* KD U-87 MG cells following TWEAK-induced non-canonical NF-κB activation. In the absence of ETS1, we unexpectedly found that almost 60% of all p52 DNA binding sites were lost (Fig. 4a) while 34 binding sites were upregulated (too few sites to be effectively seen in the figure), indicating that ETS1 regulates genome-wide p52 binding. Furthermore, approximately 60% of the p52:ETS1 co-association sites were among the lost p52 sites (Fig. 4b,c), suggesting that p52 binds indirectly to the DNA through ETS1 at the genomic loci where both TFs cooperate. We also noted that these co-localized regions only constitute a subset of all lost p52 binding sites, implying that ETS1 can also indirectly regulate and dictate p52 DNA binding through alternate mechanism(s).

**Figure 4.**
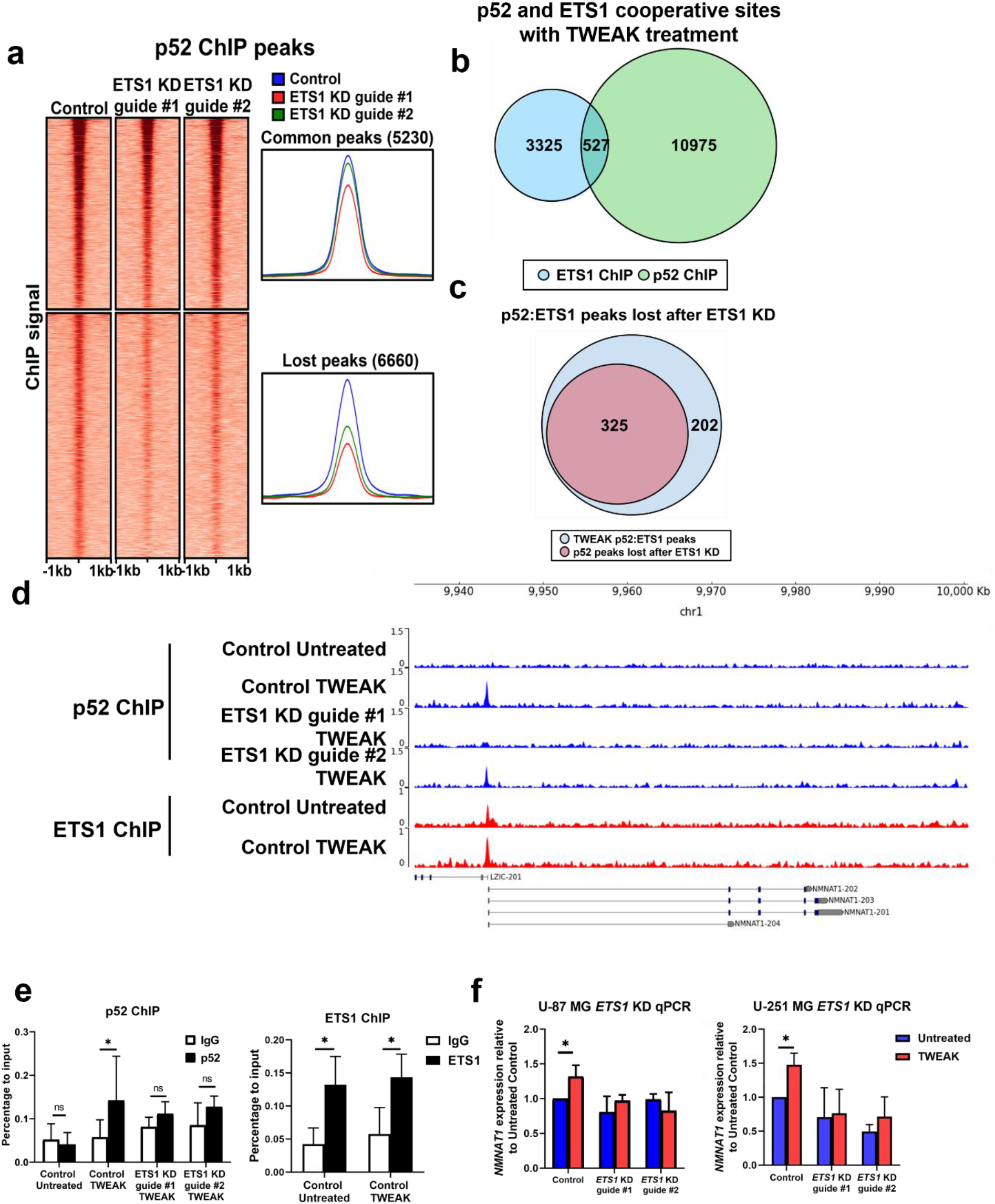
ETS1 regulates the genome-wide binding of p52. (a) DNA binding dynamics of p52 in TWEAK treated U-87 MG cells following *ETS1* KD. Venn diagram depicting (b) the number of cooperative sites between p52 and ETS1, and (c) the number of p52:ETS1 co-binding sites lost following *ETS1* KD. ChIP-seq tracks showing p52 and ETS1 binding sites following *ETS1* KD and NF-κB/p52 activation at the (d) *NMNAT1/LZIC* promoter. (e) ChIP analysis of CRISPR control and *ETS1* KD U-251 MG cells with and without, TWEAK stimulation, using p52 and ETS1 antibodies, and IgG as a negative control. (f) Relative *NMNAT1* expression was analyzed in CRISPR control and *ETS1* KD U-87 MG and U-251 MG cells, following TWEAK stimulation. Two-way ANOVA was used for all statistical analysis. \**P* < 0.05; \*\**P* < 0.01; \*\*\**P* < 0.001.

To validate the requirement of ETS1 in the maintenance of p52 binding at sites of cooperation, we selected the genomic locus of the *Nicotinamide mononucleotide adenylyltransferase 1 (NMNAT1*) gene as its promoter was bound by ETS1, independent of p52 activation. During TWEAK-induced non-canonical NF-κB activation, p52 binding was found to be enriched at the *NMNAT1* promoter. In contrast, the enrichment of p52 at the *NMNAT1* promoter was diminished when ETS1 expression was repressed (Fig. 4d). These changes were also consistent in U-251 MG cells (Fig. 4e). Further analysis also showed that the functional cooperativity between p52 and ETS1 following TWEAK treatment resulted in the upregulation of *NMNAT1* gene expression. Conversely, CRISPR-mediated downregulation of ETS1 expression, which attenuated p52 binding, abolished the induction of *NMNAT1* expression (Fig. 4f). Altogether, these results demonstrate the ability of ETS1 to dictate p52 genome-wide DNA binding through direct and indirect mechanisms. Furthermore, our data highlights the cooperative nature of p52 and ETS1, whereby the co-localization of both TFs at gene promoters can enhance target gene expression.

### p52:ETS1 regulation of *NMNAT1* drives glioma invasion and proliferation

Having identified *NMNAT1* as a target gene of p52 and ETS1 cooperativity, we proceeded to validate its functional roles in glioma progression by performing short hairpin RNA (shRNA) mediated KD of the gene in U-87 MG and U-251 MG cell lines. Here, we observed dramatic changes in the cell morphology of both glioma cell lines following downregulation of *NMNAT1* expression, which is characterised by the loss of cell membrane integrity (Supplementary Fig. 5, Fig. 5a). Further investigation showed that the loss of *NMNAT1* expression resulted in the significant downregulation of glioma cell invasion and proliferation, which was not rescued by TWEAK treatment (Fig. 5b-d). These observations confirm the oncogenic role of *NMNAT1* in glioma cells and suggest that the p52:ETS1 driven *NMNAT1* expression is crucial for glioma survival and progression. Our data further provides functional evidence for the relevance of p52:ETS1 cooperativity in the transcriptional regulation of glioma-promoting genes.

**Figure 5.**
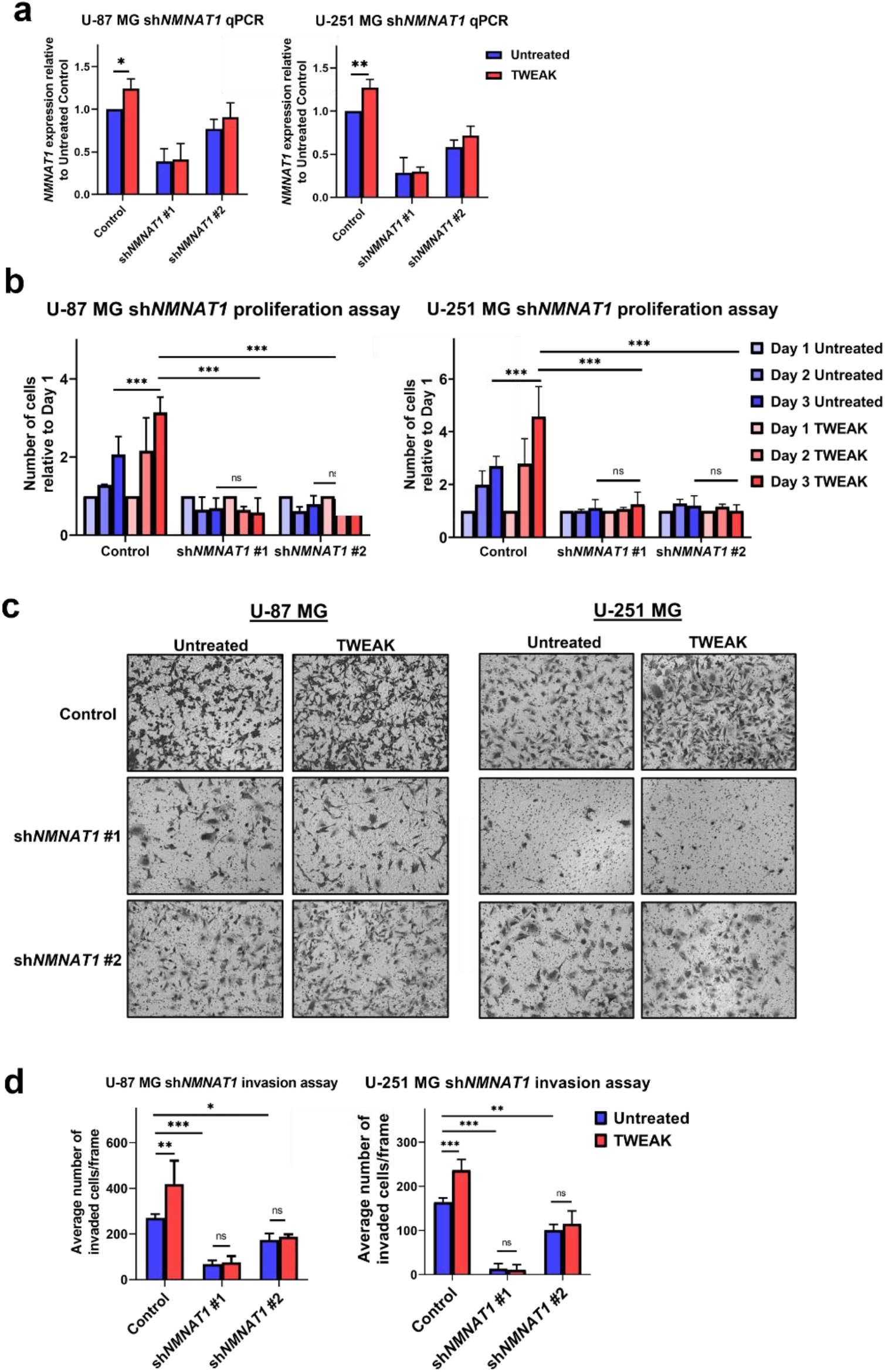
p52:ETS1 cooperativity promotes glioma cell invasion and proliferation through NMNAT1. (a) Relative *NMNAT1* expression was analysed to validate the shRNA-mediated repression of *NMNAT1* in U-87 MG and U-251 MG cells. (b) Plot depicts the average number of cells relative to day 1 (mean ± s.d) in U-87 MG and U-251 MG sh*NMNAT1*. (c) Transwell invasion assay was performed in U-87 MG and U-251 MG sh*NMNAT1* KD cells. Representative images from *n=3* biological replicates are shown. (d) Plot depicts average number of invaded cells (mean ± s.d) across five fields. All functional experiments were compared in the presence and absence of TWEAK treatment. Two-way ANOVA was used for all statistical analyses. *\*P* < 0.05; \*\**P* < 0.01; \*\*\**P* < 0.001.

## Discussion

In this study, we uncovered *ETS1* as a novel target gene of the non-canonical NF-κB pathway. The roles of non-canonical NF-κB signalling and ETS1 have both been well documented to regulate glioma migration, invasion and cell proliferation (32). Furthermore, p52 and ETS1 have been described to function cooperatively at the mutant *TERT* promoter to drive telomerase expression in gliomas (6, 18). These studies implicate the critical role of p52:ETS1 cooperativity in the transcriptional activation of cancer driver genes. Here, we provide new evidence for the direct regulation of p52 at the *ETS1* promoter to activate its expression. The overexpression of ETS1 alters its genomic and transcriptional dynamics, which in turn impact glioma invasion and survival.

On its own, ETS1 is expressed in an autoinhibited form (33) and is unable to bind to DNA. But in several cancers, it has been shown to dimerize with other TFs, such as Pax5 (34), Runx1 (31) and Runx2 (35). This increases its DNA binding affinity which can enhance the transcription of new target genes and potentially regulate oncogenic processes. Here, we report a similar signature whereby TWEAK-induced p52 activity resulted in the increased enrichment of oncogenic motifs, such as various ETS factors, KAISO and ZEB1, at ETS1 binding sites and augmented genome-wide ETS1 DNA binding which altered the transcriptomic landscape. We also observed that TWEAK-regulated ETS1 DNA binding was highly enriched in glioma patients and ETS1 activation consequently supports glioma invasion and proliferation. Altogether, these results highlight the functional significance and clinical relevance of p52-driven ETS1 in glioma-specific progression.

As reports indicate that ETS1:p52 can cooperatively activate non-κB and/or non-ETS target genes (6, 18), we investigated this regulatory mechanism genome-wide. Unexpectedly, we found that nearly 60% of all p52 binding sites were lost following ETS1 KD in TWEAK-activated glioma cells. This data suggests that ETS1 is required to sustain p52 DNA binding. We further noted that among these lost p52 sites, a small proportion comprised of the co-localized p52:ETS1 regions while almost 95% of the lost p52 peaks did not interact with ETS1 (Fig. 3b,c). This implies that ETS1 primarily regulates the p52 genomic binding landscape through indirect mechanism(s). We have earlier suggested that at sites of cooperativity, p52 binds indirectly to DNA through ETS1. Similarly, NF-κB has been reported to interact with AP-1 TFs via recruitment, without binding directly to DNA (36). As AP-1 TFs are direct target genes of ETS1 (37), it is plausible that p52-activated ETS1 enhances AP-1 expression which increases the availability of DNA-bound AP-1, thereby providing an anchor for p52 to establish indirect DNA binding.

To investigate the direct cooperation between p52 and ETS1, we focused on the genomic locus of *NMNAT1*. NMNAT1 is an enzyme involved in the NAD^+^ salvage pathway and its involvement in cancer progression has been documented in gliomas (38), osteosarcoma (39) and acute myeloid leukemia (40). We showed that in the absence of non-canonical NF-κB signalling, binding of ETS1 to the *NMNAT1* promoter is insufficient to promote its regulation. But during TWEAK induction, p52 is enriched at the *NMNAT1* promoter, resulting in p52:ETS1 co-localization. This leads to the upregulation of NMNAT1, thereby promoting glioma progression. While our data highlight the prerequisite of ETS1 for sustaining p52 DNA binding and provide genomic evidence for the indirect binding of p52 to DNA through ETS1, this study further implicates the importance of TWEAK-regulated p52:ETS1 cooperativity in the transcriptional regulation of glioma-promoting genes.

Our study highlights a previously unrecognised role of non-canonical NF-κB activation in ETS1 regulation, whereby p52-induced ETS1 expression results in a multitude of genomic, transcriptional and functional events in glioma cancers. We further investigated the direct cooperative role of TWEAK-driven p52:ETS1 and demonstrated its capacity to regulate and drive glioma progression. Intriguingly, we found that NF-κB-driven ETS1 regulated p52 DNA binding predominantly through indirect means, highlighting an overall complex regulatory relationship between these TFs. Taken together, our results emphasise the relevance of non-canonical NF-κB activation and ETS1 in glioma progression, and therefore highlights the immense potential of targeting the p52-ETS1 regulatory axis in cancer therapy.

## Methods

### Cell lines and reagents

U-87 MG and U-251 MG GBM cell lines were obtained from ATCC and tested to be mycoplasma free using the Mycoplasma PCR Detection Kit (Applied Biological Materials Inc; G238), prior to experimentation. Cells were maintained in Dulbecco’s modified Eagle’s medium (DMEM; HyClone) supplemented with 10% fetal bovine serum (FBS; Sigma). Recombinant human TWEAK (PeproTech; 310-06) was reconstituted according to the manufacturer’s recommendations. All GBM cell lines were treated with 10 ng/ml of TWEAK. Plasmids lentiCRISPRv2 (Addgene; Cat #52961) was used for generating ETS1 KD in U-87 MG and U-251 MG cell lines. sgRNAs targeting ETS1 were designed on Benchling (41), where sgRNAs with the highest On-Target score and Off-Target score were selected. sgRNAs primers used:

ETS1 guide #1 F: CACCGGGTTTCTGTCCACTGCCGG ETS1 guide #1 R: CCGGCAGTGGACAGAAACCCGGTG ETS1 guide #2 F: CACCGCTGGGCTCTGAGAACTCCGA ETS1 guide #2 R: AAACTCGGAGTTCTCAGAGCCCAGC NFKB2 guide #1 F: CACCGTGGCCCCTACCTGGTGATCG NFKB2 guide #1 R: AAACCGATCACCAGGTAGGGGCCAC NFKB2 guide #2 F: CACCGCTTTCGGCCCTTCTCACTGG NFKB2 guide #2 R: AAACCCAGTGAGAAGGGCCGAAAGC pLKO.TRC (Addgene; Cat#10878) was used for generating shNMNAT1 KD in U-87 MG and U-251 MG cell lines. shRNA primers used: shNMNAT1 #1 F: CCGGGAAGGTACACAGTTGTCAAAGCTGCAGCTTTGACAACTGTGTACCTTCTTTTTG shNMNAT1 #1 R: AATTCAAAAAGAAGGTACACAGTTGTCAAAGCTGCAGCTTTGACAACTGTGTACCTTC shNMNAT1 #2 F: CCGGGTGAATGAATGGATCGCTAATCTGCAGATTAGCGATCCATTCATTCACTTTTTG shNMNAT1 #2 R: AATTCAAAAAGTGAATGAATGGATCGCTAATCTGCAGATTAGCGATCCATTCATTCAC

### Lentivirus production and transduction

For lentivirus production, 3×10^6^ HEK293T cells were plated per 10cm plate. The next day, lentiCRISR v2 or pLKO with pMDL, VSVG and REV were transfected into the cells via calcium chloride method. The cells were washed after 8h and replenished with DMEM supplemented with 10% FBS. After 24h, supernatant containing viral particles were collected.

For lentivirus transduction, virus and polybrene were added to GBM cells. Media was changed 24h after transduction and appropriate antibiotics were added 48h after transduction.

### Cell proliferation assay

U-87 MG and U-251 MG cells were seeded at a density of 5×10^4^ and 1×10^4^ cells per well in a 12 well plate. Cells were incubated overnight to allow attachment before being starved for 8h in serum-free DMEM. After which, the number of cells were counted (this number will be referred to as day 1) and 10ng/ml TWEAK was added. After 24h and 48h, the number of cells were counted again.

### Invasion assay

Invasion assay was performed using 24 well Transwell inserts (Corning). Matrigel (Corning) was diluted to 200ug/ml with Optimem (Gibco), 100ul was added into each insert and incubated at 37C for 2h. GBM cells were starved for 8h beforehand, subsequently, the GBM cells were treated with 10ng/ml TWEAK. After 24h, the GBM cells were trypsinized, counted, resuspended in 100ul optimem containing 10ng/ml TWEAK and seeded above the Matrigel layer. 600ul DMEM supplemented with 10% FBS was added to the lower chamber. The GBM cells were then incubated at 37C for 24h. Following which, the transwell inserts were fixed in 70% ethanol, stained in 0.2% crystal violet and non-invading cells in the top chamber were removed with cotton buds. Invading cells were then visualised with an inverted microscope at 5 different frames. The number of invading cells was counted in each frame and averaged.

### Western blot analysis

Western blotting was performed using standard methods and the following antibodies were used for analysis: anti-NFkB2 (Cell Signalling; 3017), anti-ETS1 (Cell Signalling; 14069) and anti-GAPDH (Santa Cruz; 32233). Antibody dilutions used were 1:10000 for anti-GAPDH and 1:1000 for the other antibodies.

### RNA extraction and quantitative real-time PCR (qPCR)

Total RNA was isolated using TRIzol Reagent (Invitrogen) according to the manufacturer’s protocol. Complementary DNA was synthesized using reverTra Ace qPCR RT Master mix with gDNA remover (Toyobo; FSQ-301) and qPCR was performed in triplicates using iTaq Universal SYBR Green Supermix (Bio-Rad; 1725125). The sequences of the qPCR primers used are: NMNAT1 F: TCACCAACATGCACCTCAGG NMNAT1 R: AGTTCTGCCATGATGACCCG GAPDH F: GCATCCTGGGCTACACTGA GAPDH R: CCACCACCCTGTTGCTGTA

### RNA library construction, sequencing and analysis

Total RNA was treated with TURBO DNA-free kit (Invitrogen; AM1907). RNA-seq library was constructed with NEBNext Ultra II Directional RNA Library Prep Kit for Illumina (NEB; E7765) through the polyA mRNA workflow. RNA-seq libraries were sequenced on HiSeqX. All RNA-seq experiments were performed with three biological replicates.

The quality of sequencing reads was assessed using FastQC (42) and adaptors were removed with Trimmomatic (43). RNA-seq reads were mapped to hg38 with STAR (44) and processed by featureCounts (45). Differential analysis was performed with DESeq2 (46) where differentially expressed genes were defined by having p value ≤ 0.05 and fold change ≥ 1.5. Gene ontological analysis was performed using PANTHER (47, 48).

### Chromatin immunoprecipitation (ChIP), library construction, sequencing, and analysis

ChIP was performed on GBM cell lines as described previously (6), using the following antibodies: anti-NFkB2 (Bethyl Laboratories; A300-BL7039), anti-ETS1 (Active Motif; 39580), anti-RNA polymerase II clone CTD4H8 (EMD Millipore; 05-623) and non-specific rabbit IgG (Bethyl Laboratories; P120-101). Eluted DNA was used for qPCR or sequencing library preparation with NEBNext Ultra II DNA Library Prep Kit (NEB; E7645) according to the manufacturer’s protocol.

qPCR was performed using Sso advanced Universal SYBR mix (Bio-rad; 1725272). Ct values were normalized to relative amount of 1% of input DNA. The sequences of the ChIP qPCR primers used are: NMNAT1 promoter F: CATGGTCCCCGAAACTCCTT NMNAT1 promoter R: CTAAACTCTGGGACTGGCGG

ChIP DNA libraries were sequenced on HiSeqX. The quality of sequencing reads were assessed using FastQC (42). Trimmomatic (43) was used to remove adaptors, and verified by running FastQC once more. Reads were mapped to hg38 using Bowtie2 (49) and peaks were called using MACS2 with subtraction of input signal. Blacklisted regions were removed from the called peaks using bedtools (50). The gained and lost ETS1 peaks were determined using bedtools. Differential RNA polymerase II peaks were analysed using DiffBind (51).

Heatmaps showing signal distribution over ChIP peak regions were generated using deepTools (52). ETS1 binding motifs were identified by HOMER (53) with the hg38 genome as background and MEME-ChIP (54).

### Integration with TCGA pan-cancer RNA-seq and ATAC-seq data

Pan-cancer patient RNA-seq and ATAC-seq sample counts (25) from The Cancer Genome Atlas (TCGA) were retrieved using the UCSC Cancer Browser (http://xena.ucsc.edu/welcome-to-ucsc-xena/) (55). Heatmaps were generated using R Statistical Programming Language and ggplot2 (56).

### Statistical analysis

In all figures, the figure legends denote the statistical tests used, kind of replicates and the value of *n*. Asterisks represent the degree of statistical significance as described in the figure legends. All statistical analyses and graphics were performed using GraphPad Prism software.

## Acknowledgements

This work is supported by the National Research Foundation (NRF) Singapore, under the Singapore NRF Fellowship (NRF-NRFF2018-04). We thank the Nanyang Technological University for the PhD scholarship support of N.S. and Nanyang Assistant Professorship start-up of Y.L.’s lab.

## Author contributions

Y.L. conceptualized the ideas for this manuscript and supervised the project. N.S. and Y.L. planned and devised the experiments. N.S. performed all experiments and computational analyses. N.S. wrote the manuscript and Y.L. edited it.

## Competing interests

The authors declare no competing interests.

## Data availability statement

The datasets generated during and/or analysed during the current study will be uploaded onto the NCBI Sequence Read Archive (SRA) repository before publication.

## Notes

There is no conflict of interest.

http://xena.ucsc.edu/welcome-to-ucsc-xena/

### Competing Interest Statement

The authors have declared no competing interest.

