## Supplementary Figures for "NF-κB/p52 augments ETS1 binding genome-wide to promote glioma progression"

**Figure S1**

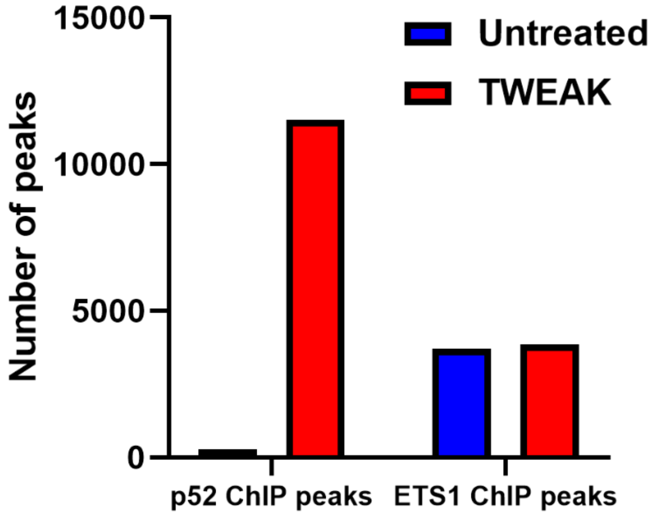


**Supplementary Figure 1. Number of p52 and ETS1 ChIP peaks in U-87 MG cells with and without TWEAK treatment.**

**Figure S2**

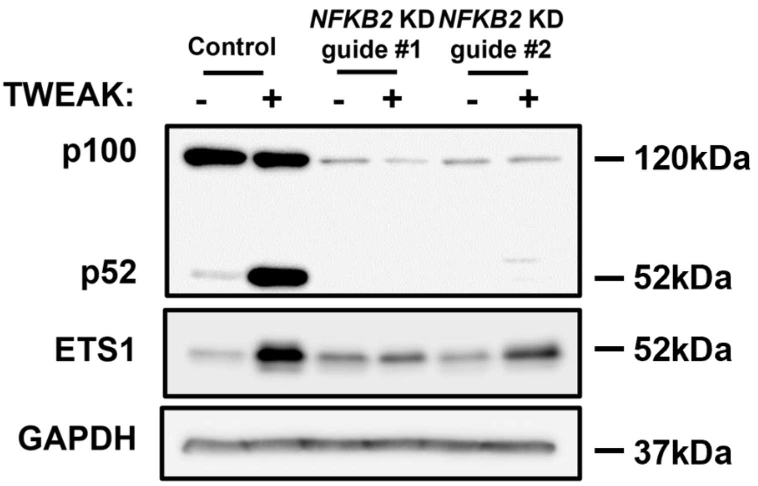


**Supplementary Figure 2. p52 and ETS1 expression in U-87 MG cells following NFKB2 knockdown and TWEAK treatment analysed through western blotting.**

### Figure S3

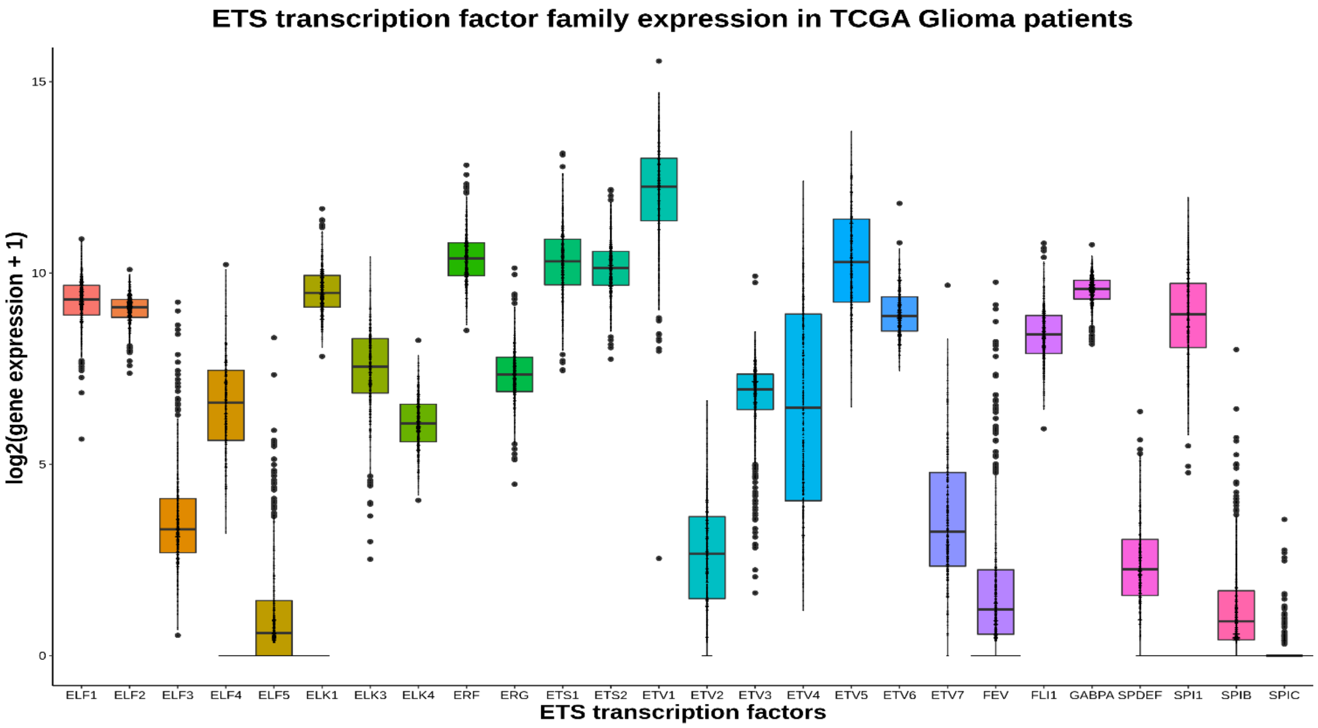


**Supplementary Figure 3. Gene expression of ETS transcription factors in glioma patients (*n* = 702) from TCGA RNA-seq dataset.**

### Figure S4

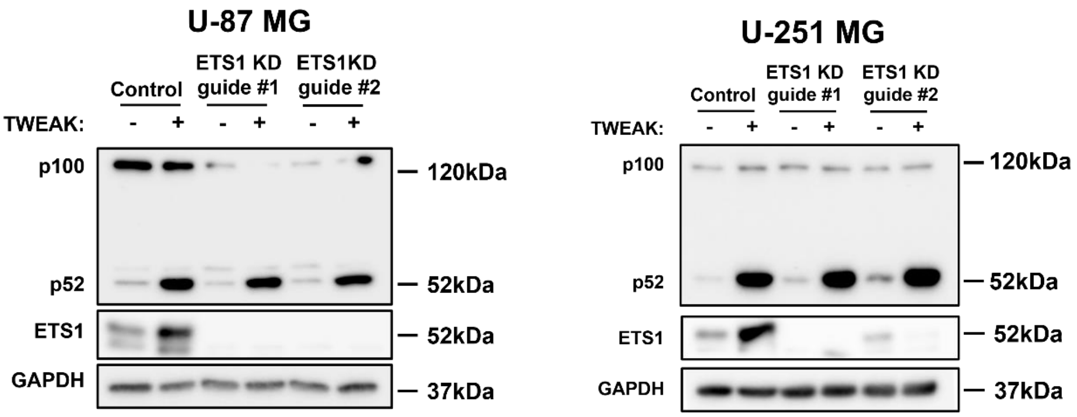


**Supplementary Figure 4. p52 and ETS1 expression in U-87 MG and U-251 MG cells following *ETS1* knockdown and TWEAK treatment analysed through western blotting.**

**Figure S5**

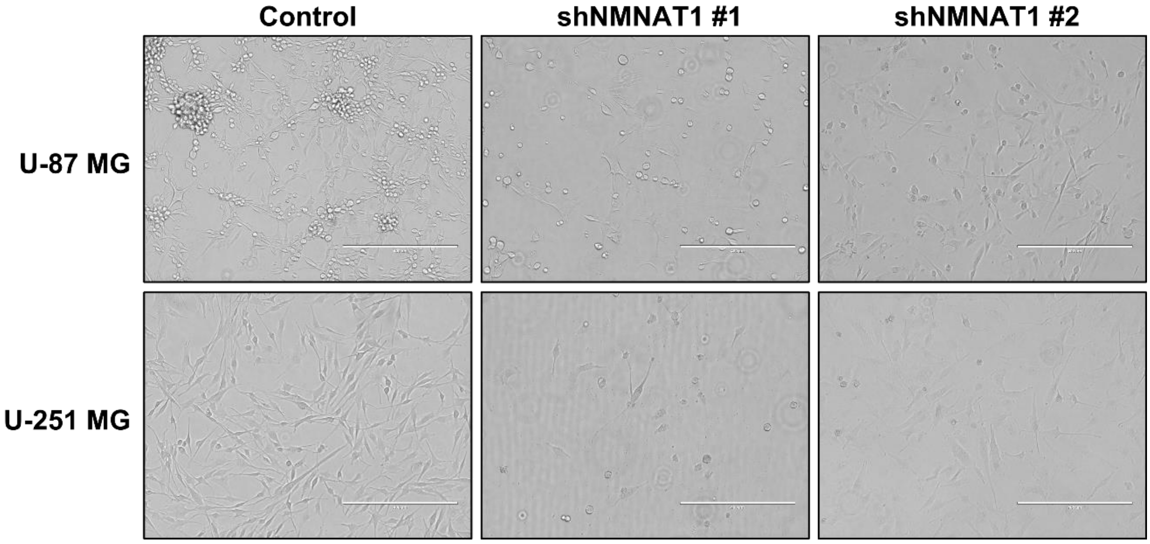


**Supplementary Figure 5. Cell morphology of U-87 MG and U-251 MG cells following shRNA-mediated NMNAT1 repression.**
